# Spatiotemporal Mapping of Brain Cilia Reveals Region-Specific Oscillation of Length and Orientation

**DOI:** 10.1101/2023.06.28.546950

**Authors:** Roudabeh Vakil Monfared, Sherif Abdelkarim, Pieter Derdeyn, Kiki Chen, Hanting Wu, Kenneth Leong, Tiffany Chang, Justine Lee, Sara Versales, Surya M. Nauli, Kevin Beier, Pierre Baldi, Amal Alachkar

**Author notes:** These authors contributed equally to this work (Roudabeh Vakil Monfared, Sherif Abdelkarim). **Corresponding Authors:** Amal Alachkar Department of Pharmaceutical Sciences University of California, Irvine 356A Med Surge II Irvine CA, 92697-4625; Pierre Baldi, PhD Departments of Computer Science School of Information and Computer Sciences University of California, Irvine Irvine CA, 92697.

## Abstract

In this study, we conducted high-throughput spatiotemporal analysis of primary cilia length and orientation across 22 mouse brain regions. We developed automated image analysis algorithms, which enabled us to examine over 10 million individual cilia, generating the largest spatiotemporal atlas of cilia. We found that cilia length and orientation display substantial variations across different brain regions and exhibit fluctuations over a 24-hour period, with region-specific peaks during light-dark phases. Our analysis revealed unique orientation patterns of cilia at 45° intervals, suggesting that cilia orientation within the brain is not random but follows specific patterns. Using BioCycle, we identified circadian rhythms of cilia length in five brain regions: nucleus accumbens core, somatosensory cortex, and three hypothalamic nuclei. Our findings present novel insights into the complex relationship between cilia dynamics, circadian rhythms, and brain function, highlighting cilia crucial role in the brain’s response to environmental changes and regulation of time-dependent physiological processes.

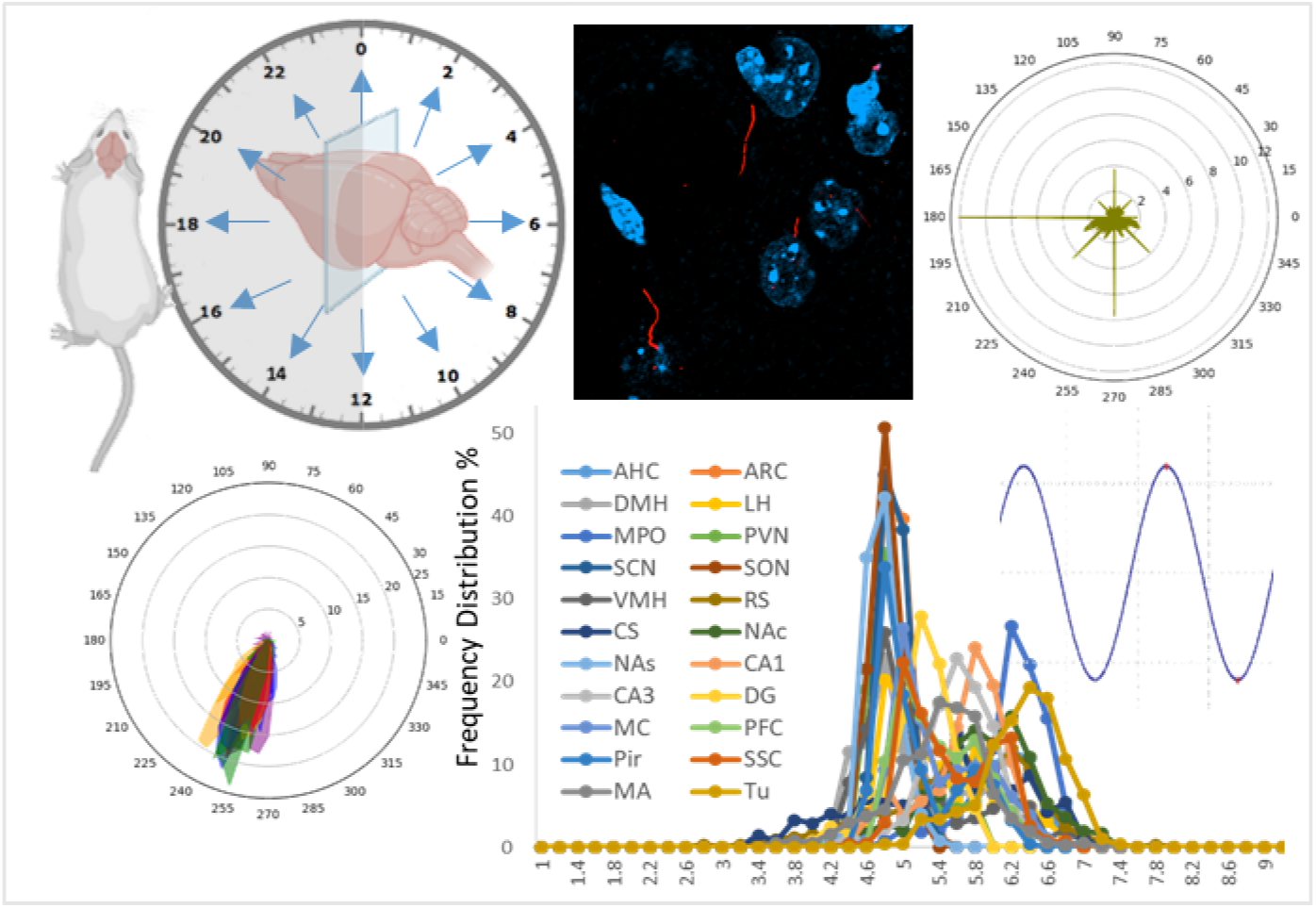

## Introduction

Cilia are evolutionarily-conserved, microtubule-based organelles that extend from the surface of eukaryotic cells, including neurons in the mammalian brain. Their presence in diverse organisms from single-celled organisms to complex multicellular animals emphasizes their fundamental importance in a wide range of biological processes. Primary cilia, in particular, serve as essential sensory and signaling hubs, playing critical roles in cellular homeostasis, development, and signal transduction [1-9].

A striking feature of cilia is their dynamic nature, which is manifested in the continuous modulation of their structure and function in response to various environmental cues and cellular signaling events [8-16]. Mechanical forces or cerebrospinal fluid (CSF) flow can induce elongation or shortening of cilia, while intracellular signaling pathways mediated by cyclic AMP (cAMP) or calcium can reversibly modulate their morphology [17-20]. We and others have shown that neurotransmitters such as dopamine, serotonin, melanin concentrating hormone (MCH), and adenosine can modulate primary cilia length, affecting their signaling capacity [18, 21-24]. Although primary cilia are non-motile, their orientation can be dynamic due to the underlying cellular structures and signaling pathways regulating their relative positions. This dynamic orientation allows primary cilia to adapt to environmental changes and respond to different extracellular cues more effectively. Cilia orientation may adjust in response to numerous and varied cellular events, such as during cell migration [25, 26], tissue repair [25], or in response to extracellular stimuli [27] such as mechanical stress or chemical gradients.

Many environmental stimuli and cellular signaling events sensed by cilia often follow rhythmic temporal patterns, such as light--dark cycles and fluctuations in temperature or nutrient availability [28-34]. These patterns also extend to a broad range of functions influenced by cilia signaling, such as daily sleep-wake cycles, feeding behaviors, sexual/reproductive behaviors, energy balance, metabolism, and regulation of body temperature [35-42]. Furthermore, G-protein coupled receptors (GPCRs) located on primary cilia, including MCHR1, MC4R, DRD1, DRD2, and GALR, are involved in various circadian functions such as food intake, body temperature regulation, and sleep [37, 43-47]. We have previously uncovered a high degree of circadian rhythmicity and spatiotemporal oscillations in cilia-associated gene expression across primate brain areas, together with dynamic shifts in the expression of genes associated with primary cilia throughout the human lifespan [48, 49]. This was further supported by our research into cilia gene dysregulation in major psychiatric disorders, hinting at shared pathophysiological processes [50]. This implies that cilia signaling dysfunctions may underlie common neurological deficits across these disorders.

Despite the considerable advancements in our knowledge about the structural dynamics of primary cilia, there is currently very little evidence for such cilia dynamics and circadian rhythms. Thus, the main aim of this study is to study the dynamic nature of primary cilia in the mouse brain by characterizing the spatiotemporal variations in cilia length and orientation, and investigating their circadian patterns. This comprehensive analysis of cilia properties and their circadian dynamics will provide a valuable framework for future studies to shed light on the functional roles of cilia in different brain regions and their contribution to the fine-tuning of circadian functions and other neurological processes.

## Methods

### Animals

Eight-week-old Swiss Webster male mice were obtained from Charles River (North Carolina, USA). The mice were maintained on a 12-hour light/dark cycle and provided with food and water ad libitum. All experimental procedures were approved by the Institutional Animal Care and Use Committee (IACUC) of the University of California, Irvine, and were conducted in accordance with national and institutional guidelines for the care and use of laboratory animals.

### Brain Tissue Collection

Mice were deeply anesthetized using isoflurane and subsequently transcardially perfused with a combination of 0.9% saline and 4% paraformaldehyde (PFA) (Ref). We employed a total of 56 mice, perfusing 4-5 mice at every two-hour interval over a 24-hour period, resulting in 12 time-points. Following perfusion, we harvested the brains, preserved them in PFA overnight, and then transferred them to a 30% sucrose solution. The brains were then coronally sectioned at a thickness of 30 μm using a microtome.

### Immunohistochemistry and Imaging

For immunohistochemistry preparation, 3-4 sections were selected from various levels across the brain. The sections were then blocked using goat serum in PBS containing 0.3% Triton X-100 for 1 hour. Following this step, the brain sections were incubated in blocking buffer solution containing the primary antibody, ADCY3 (LSBIO-C204505 ADCY3), at a dilution of 1:500. Post primary antibody incubation, the sections were rinsed with PBS and incubated with the secondary antibody, Invitrogen AlexaFluor546 goat anti-rabbit, at a dilution of 1:500, along with DAPI at a dilution of 1:10,000 (Thermo Fisher Scientific). Subsequently, the sections were washed with PBS and mounted onto slides. The cilia were imaged using the Keyence BZ-9000 fluorescence microscope with a 20x objective lens. Brain regions analyzed include: Anterior hypothalamic area (AHC), Arcuate nucleus (ARC), Dorsomedial hypothalamus (DMH), Lateral hypothalamus (LH), Median preoptic area (MPO), Hypothalamic paraventricular nucleus (PVN), Suprachiasmatic nucleus (SCN), Supraoptic nucleus (SON), Ventromedial hypothalamus (VMH), Caudal striatum (CS), Nucleus accumbens core (NAc), Nucleus accumbens shell (NAs), Rostral striatum (RS), Olfactory tubercle (TU), Cornu Ammonis 1 (CA1), Cornu Ammonis 3 (CA3), Dentate Gyrus (DG), Motor cortex (MC), Prefrontal cortex (PFC), Piriform cortex (Pir), Somatosensory cortex (SSC), Medial amygdala (MA).

### Cilia Length and Angle Measurement

Our first step involved pre-processing the images by converting them to grayscale, to make them conducive for analysis and to mitigate color-related complications. We then applied Gaussian blur to the images to minimize noise or minor details that might compromise the accuracy of our measurements. Next, we used canny edge detection to find all the edges in the images. This allowed us to identify the cilia more easily and accurately. We applied a threshold value to the canny edge detection output to obtain a binary image where the edges were represented by white pixels and the background is represented by black pixels. After obtaining the binary image, we used the findContours() function in OpenCV to detect all the cilia in the image. This function identifies and segments all the distinct objects in the image, in this case the cilia, and returns a list of their contours. However, some of these contours may be noise or not relevant to our analysis. Therefore, we filtered them based on their size, aspect ratio, and other factors to only capture the ones that are likely to be cilia. For instance, we filter the contours based on their length, as cilia are typically longer than they are wide. We set a minimum and maximum length threshold for the contours to keep only the ones that fall within this range. In our implementation, we use a minimum length of 1 micron and a maximum length of 15 µm. Furthermore, we filter the contours based on their aspect ratio, as cilia are generally slender structures with a length-to-width ratio greater than 3:2. We set a minimum aspect ratio threshold for the contours to remove any that do not meet this criterion. In our implementation, we used a minimum aspect ratio of 1.5. For each contour, we fit a bounding box around it using the cv2.boundingRect() function. This gave us the length and angle of the cilia. The length was calculated as the distance between the top and bottom points of the bounding box, and the angle was calculated as the angle between the major axis of the box and the horizontal axis. However, we also needed to determine the base of the cilia accurately. To achieve this, we used the cv2.minMaxLoc() function to find the brightest pixel in the grayscale image within the bounding box of the cilia. The cilia’s base is typically the most illuminated section, and this method enables us to pinpoint it with precision. To further improve the accuracy of our angle measurements, we combine the location of the base of the cilia with the center point of the bounding box. By comparing the relative positions of these points, we were able to determine which quartile the cilia were facing and adjust the angle measurement accordingly. Finally, we saved all the information about the cilia, including their lengths, angles, and positions, in a CSV file. This allowed us to analyze the data more easily and systematically. The two programs we developed and used are provided as supplementary materials, referred to as Program 1 and Program 2.

### Circular Angle Mean Calculation

Due to the circular nature of angles, the traditional arithmetic mean method is not applicable for calculating angle means. Therefore, we computed the circular mean, also known as the angular mean, to calculate meaningful statistics for our angle dataset. The circular mean of our angle dataset was calculated, using the pandas library in Python as follows: First, the angle values were converted to radians, and all the data were subsequently transformed into sines and cosines. The means of the sines and cosines were calculated by summing the values and dividing them by the total count of each value. The circular mean was determined using the atan2 function, a variant of the arctangent function that accepts two arguments and yields results within the appropriate quadrant. The obtained results were then converted back to degrees using the equation: Degrees = radians x (180/π). If negative outputs were obtained from the atan2 function and the data ranged from 0 to 360, 360 was added to those values to align them within the correct range.

### Circadian Rhythm Analysis

BIO_CYCLE, a deep-learning model designed for assessing periodicity of signals, was utilized to determine significant circadian patterns in cilia length and angle, as well as their oscillation’s amplitude and phase [51, 52]. BIO_CYCLE is trained using synthetic and real biological time series datasets with labeled periodic and aperiodic signals. This tool comprises a deep neural network (DNN) for classification, which distinguishes periodic from non-periodic signals, and another DNN for regression estimating the signal’s period, phase, and amplitude. The oscillation status of a parameter is defined based on a p-value (with a cutoff of 0.05) computed by BIO_CYCLE as follows: N aperiodic signals are first generated from the synthetic time series datasets, and the N output values V(i) (i=1,…,N) of the classification DNN on these aperiodic signals are calculated. The N output values of the classification DNN on these aperiodic signals are then calculated, forming the basis for the null hypothesis distribution. The output value V of the new signal s is then compared to V(i)(i=1,…,N), producing the estimate for the probability of obtaining an output of size V or greater (p-value), assuming that the signal is part of the null distribution (aperiodic signals). A smaller p-value suggests a higher likelihood of the signal being periodic. The related q-values are calculated via the Benjamini and Hochberg procedure. BIO_CYCLE is freely accessible from the CircadiOmics web portal: http://circadiomics.igb.uci.edu.

### Correlation between Circadian Variables and Cilia Length and Angle

We conducted a more comprehensive examination of the relationship between cilia length, angle data, and various other circadian variables, using digitalized data from multiple studies, with permission. This digitalized data encompassed a range of circadian variables and behaviors including food intake [53], locomotor activity [54], core body temperature [55], efflux via the blood-brain barrier [56], melatonin levels [54], and EEG results like NREM and REM data [57]. The data were digitalized from graphs of wild-type subjects using the DigitizeIt software (www.digitizeit.xyz). After aligning this digitized data with our cilia length and angle information at various time intervals, we conducted correlation analyses using GraphPad Prism. For regions that showed a correlation with these variables, we performed linear regression tests, determining the correlation coefficient (r) and p-value.

### Network Analysis

Using the Allen Mouse Brain Connectivity Atlas data [58], we built a structural network of the regions used in our study. We normalized projections from source regions to other regions of interest by dividing the projection density by the injection volume. The normalized projection density was averaged across all experiments for each pair of source and target regions to generate a connectivity matrix. These values were log normalized to produce a more effective distribution for use as edge weights in a network. Community detection was carried out using the Louvain algorithm implemented in scikit-network with a resolution parameter of 1 [59]. We identified three main clusters of brain regions: purple (NAc, NAs, and others), green (PFC, MC, and others), and red (CA1, CA3, and DG). We examined the correlation between cilia length and angle with various network features. We examined all pairs of regions within and across different clusters, identifying varying fractions of region pairs correlated by either cilia length or angle. We also analyzed whether these fractions would likely occur if regions were correlated randomly, shuffling all correlations across region pairs for this purpose. We constructed cilia correlation networks using a log scaled R value to achieve a desirable distribution for edge weights. Network analysis was performed in Python using NetworkX [60], with network visualizations created using Plotly [61].

### Data Analysis

Data were Analyzed with GraphPad Prism 9.4 and was considered statistically significant if p<0.05. Data on the graph are expressed as mean ± standard error of the mean (SEM). Statistical analyses were performed to compare the means of lengths and angles at different time points using one way ANOVA. Correction for multiple comparisons was performed by controlling the False Discovery Rate (FDR), using two-stage linear step-up procedure of Benjamini, Krieger, and Yekutieli. We computed the Pearson correlation coefficient (r) for every pair of region-specific cilia lengths and angles across 24-hour time points. In addition, we generated a correlation matrix based on the correlation of cilia lengths/angles among all brain regions.

## Results

To conduct a comprehensive analysis of the spatiotemporal dynamics of cilia length and orientation across various mouse brain regions, we employed a specialized program to assess the length and angle of cilia in brain sections from an extensive sample size of over 50 mice (at least four mice per time point) (Fig. 1a, S1). We carried out sampling at 2-hour intervals throughout a 24-hour period, analyzing 3-4 sections from each mouse brain. Our analysis focused on 22 brain regions that were visibly rich in cilia (Fig. 1b). A comprehensive examination of over 10 million cilia, including data from regions containing cilia counts ranging from 153,448 (in RS) to 977,129 (in DMH), allowed us to obtain a detailed characterization of cilia in these structures of the brain (Fig. 1c).

**Figure 1:**
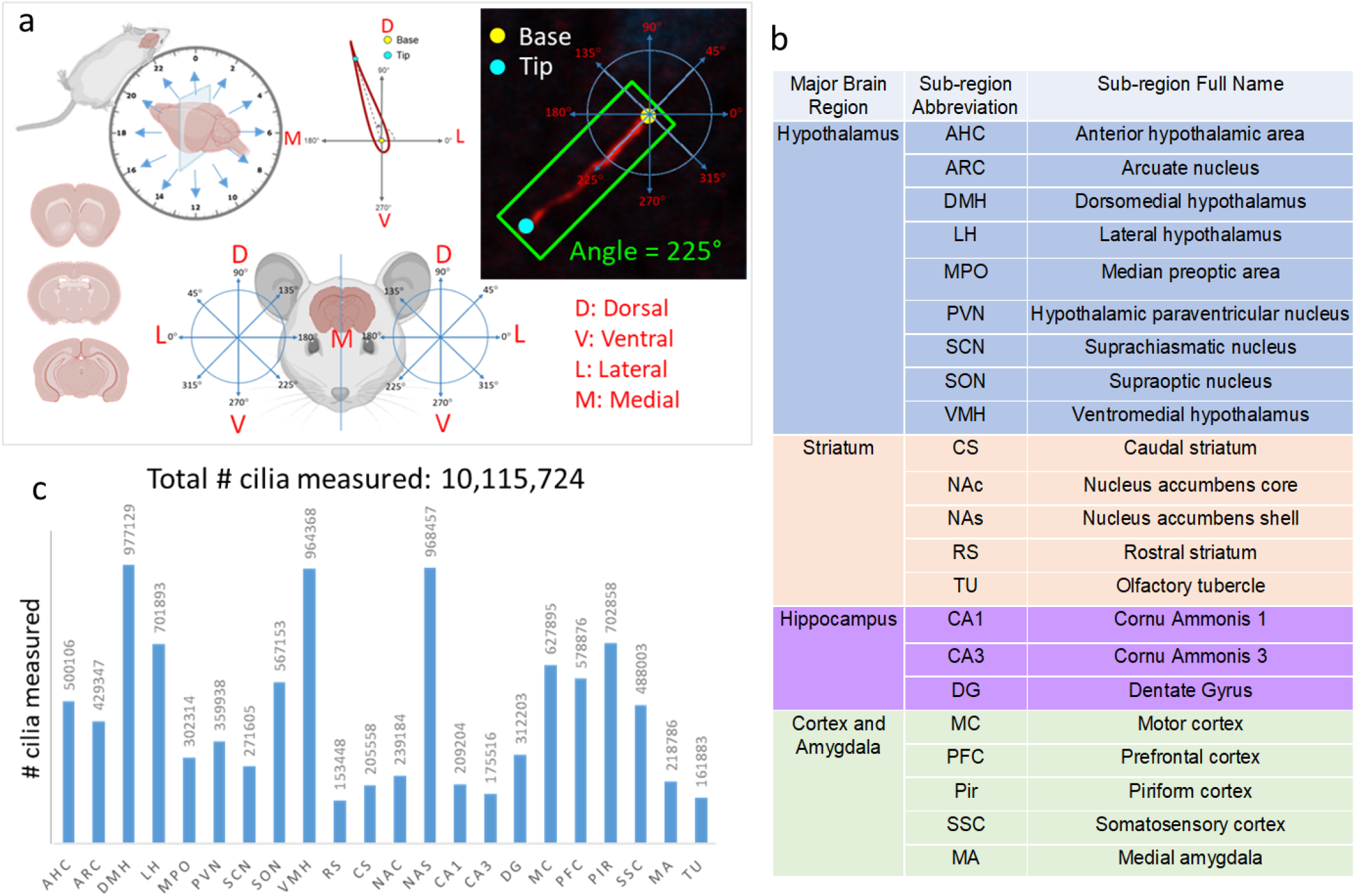
Experimental design and data analysis. **(a)** Schematic Representation: (Top Left) Mice were perfused and harvested at two-hour intervals over a 24-hour period. The harvested mice were then sectioned coronally. (Bottom Left) Coronal brain sections at different brain levels. (Top Middle) The ciliary base was defined as the origin of the coordinate system, and the tip of the cilia was labeled. Cilia length was measured from the base to the tips, and the circular angle was determined. (Bottom Middle) The brain section of the mice was divided into left and right hemispheres. For cilia found in the right hemisphere, a counter-clockwise coordinate system was used to determine the circular angle (Right). For cilia found in the left hemisphere, a clockwise coordinate system was used (Left). (Top Right) An example of the angle measurement from cilia found in the right hemisphere. **(b)** Brain Regions analyzed for cilia length and angle**. (c)** The number of cilia analyzed in 22 different brain regions of the mice.

We presented the data using two approaches. We first analyzed the lengths and angles of individual cilia across all samples to provide an in-depth view of their frequency distribution. We also calculated the mean length and angle of cilia in each section of the left and right hemisphere as a single value, which was then used in further analyses.

### Region-dependent variations in cilia length and angle in the mouse brain

Our analysis of cilia length in the whole brain and across brain regions revealed that the mean cilia length for the whole brain, was 5.15 µm, with a range across brain regions from 4.79 to 6.33 in the NAs and TU, respectively (Fig. S2a). Individual cilia lengths of 3 or 4 µm were observed as the most frequent across all brain regions examined (Fig. S2b,c). When we considered the mean cilia length from each section of the left and right hemisphere as a single value, the overall brain cilia length averaged at 5.4 ± 0.01 µm. The lengths varied across different regions, ranging from 4.87 µm in the NAs to 6.3 µm in the TU (Fig. 2a, Table S1). Except for TU, the average cilia length in all other regions was below 6 µm (Fig. 2a). In terms of frequency distribution, we found that around 40% of cilia in all regions fell within the 4.8-5.4 µm range (Fig. 2b,c, S2d).

**Figure 2:**
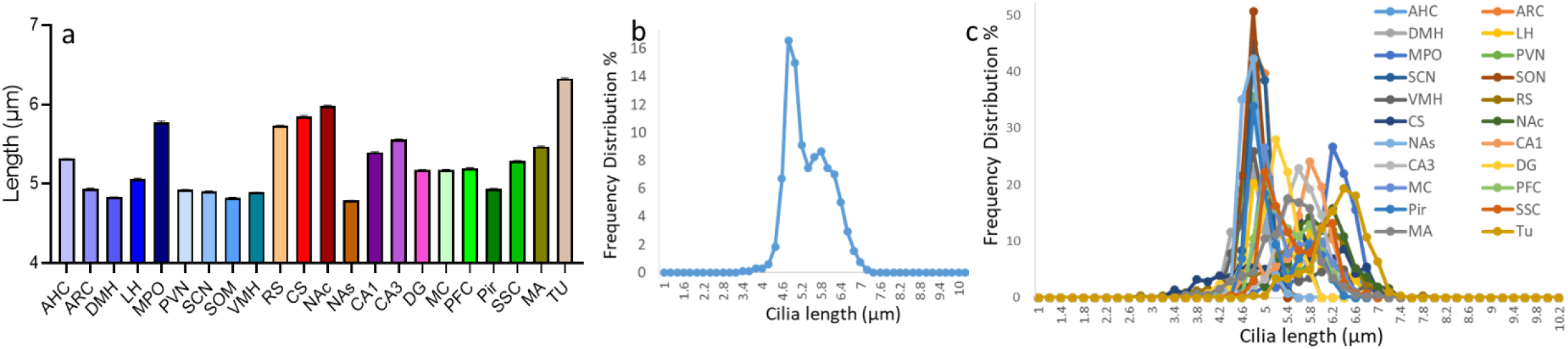
Cilia length analysis in different brain regions. **(a)** Histogram showing the mean of cilia length in the 22 brain regions based on the average length measured in each brain section. The x-axis represents cilia length in micrometers and the y-axis represents the percentage frequency of occurrence. **(b) & (c)** Histogram showing frequency distribution of cilia length in **(b)** the whole brain and **(c)** the individual 22 brain regions, based on the average length measured in each brain section. Each brain section’s measurements are consolidated into a single data point representing the respective brain region. The mean cilia length is determined by considering the average lengths from both the left and right hemisphere sections (3-4 brain sections from each region). The x-axis represents cilia length in micrometers and the y-axis represents the percentage frequency of occurrence.

Our analysis of the frequency distribution of individual cilia angles across over 10 million cilia uncovered an intriguing pattern common across all regions - the angles at which the highest frequencies occur coincide with the primary directions on a compass or Cartesian axes (0°, 45°, 90°, 135°, 180°, 225°, 270°, and 315°), but not values in between (Fig. 3a,b). Remarkably, the largest proportion of cilia were predominantly oriented towards either 180° (165-195°) or 270° (255-285°), suggesting a predominant medial and/or ventral orientation of cilia (Fig. 3a,b) when considering all samples taken across all time points in the circadian cycle. Further examination revealed that cilia orientation is not uniformly distributed across different brain regions, and that specific regions may exhibit a unique preferred angle of orientation (Fig. 3c-e). For instance, the RS, CS, MA, and CA3 displayed a predominant cilia orientation towards 180°, with a secondary preference for 270°. Conversely, regions such as the NAc, TU, DG, and CA1, showed a different pattern with a primary cilia orientation towards 270 degrees, closely followed by 180 degrees (Fig. 3e). Interestingly, we observed a more scattered distribution of cilia orientation in the hypothalamic nuclei, various cortices, and the NAs. Regardless, in all brain regions studied, cilia orientation largely followed 45°-interval patterns, with a notable dominance of medial and/or ventral orientations (Fig. 3e,f).

**Figure 3:**
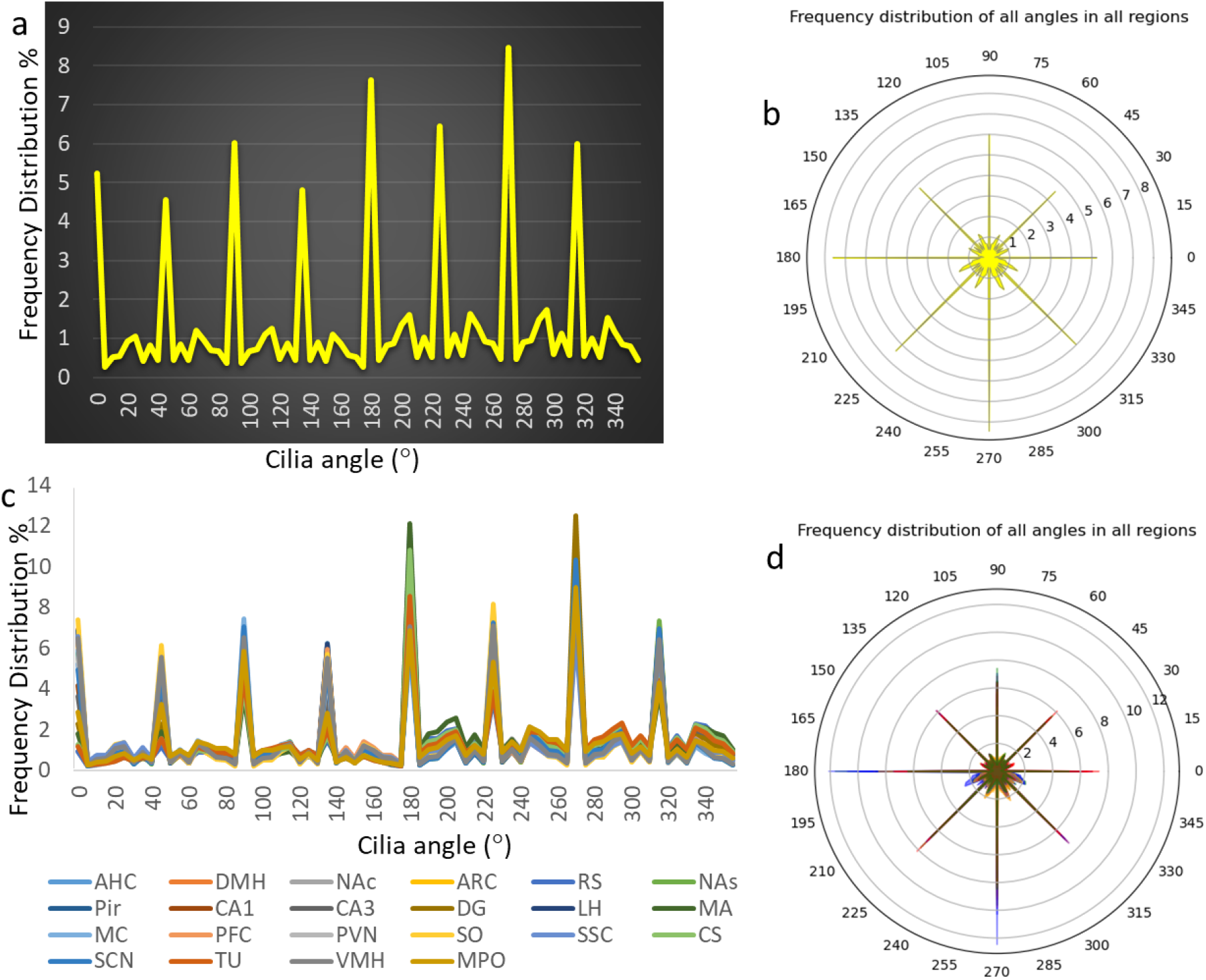
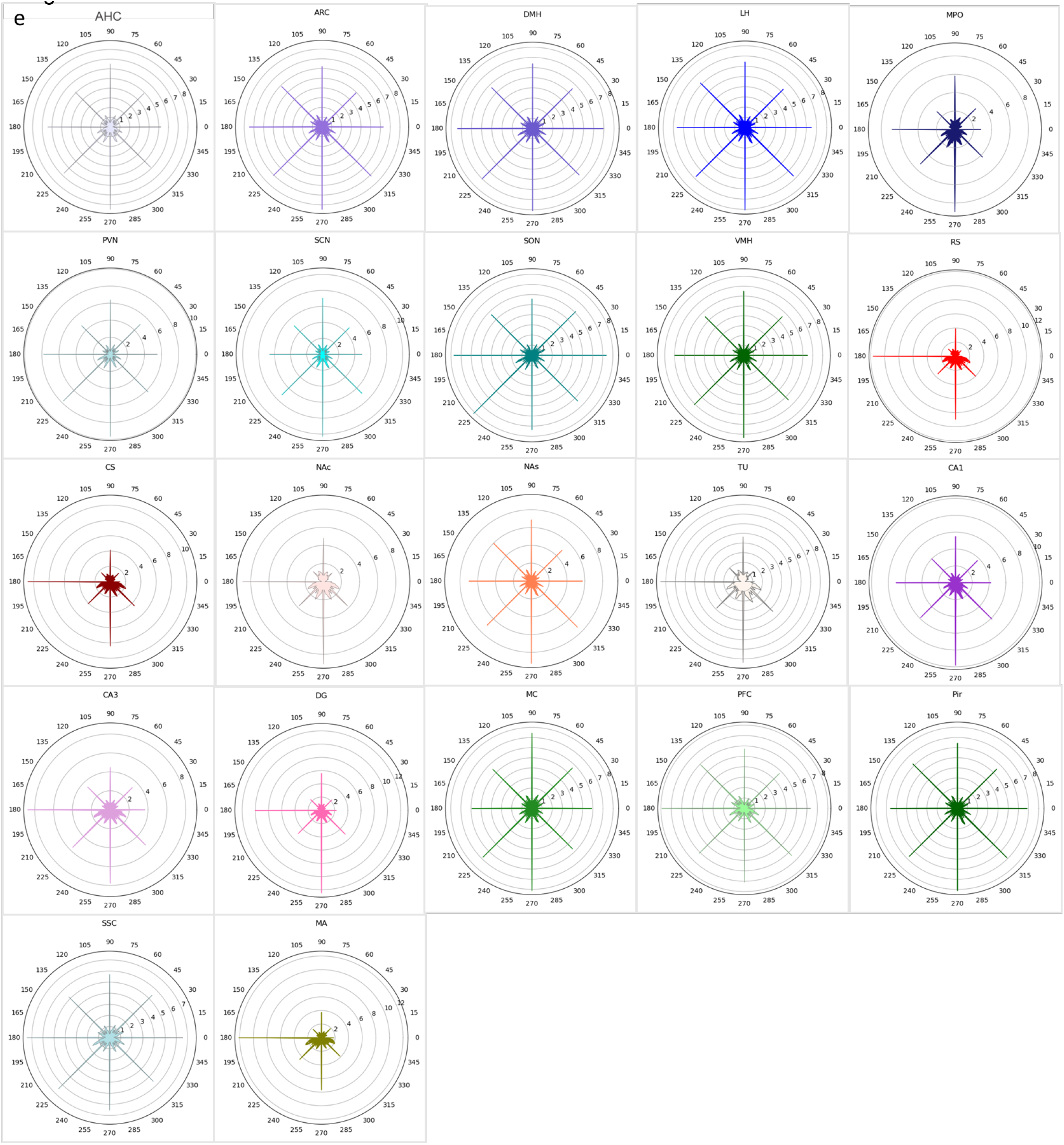
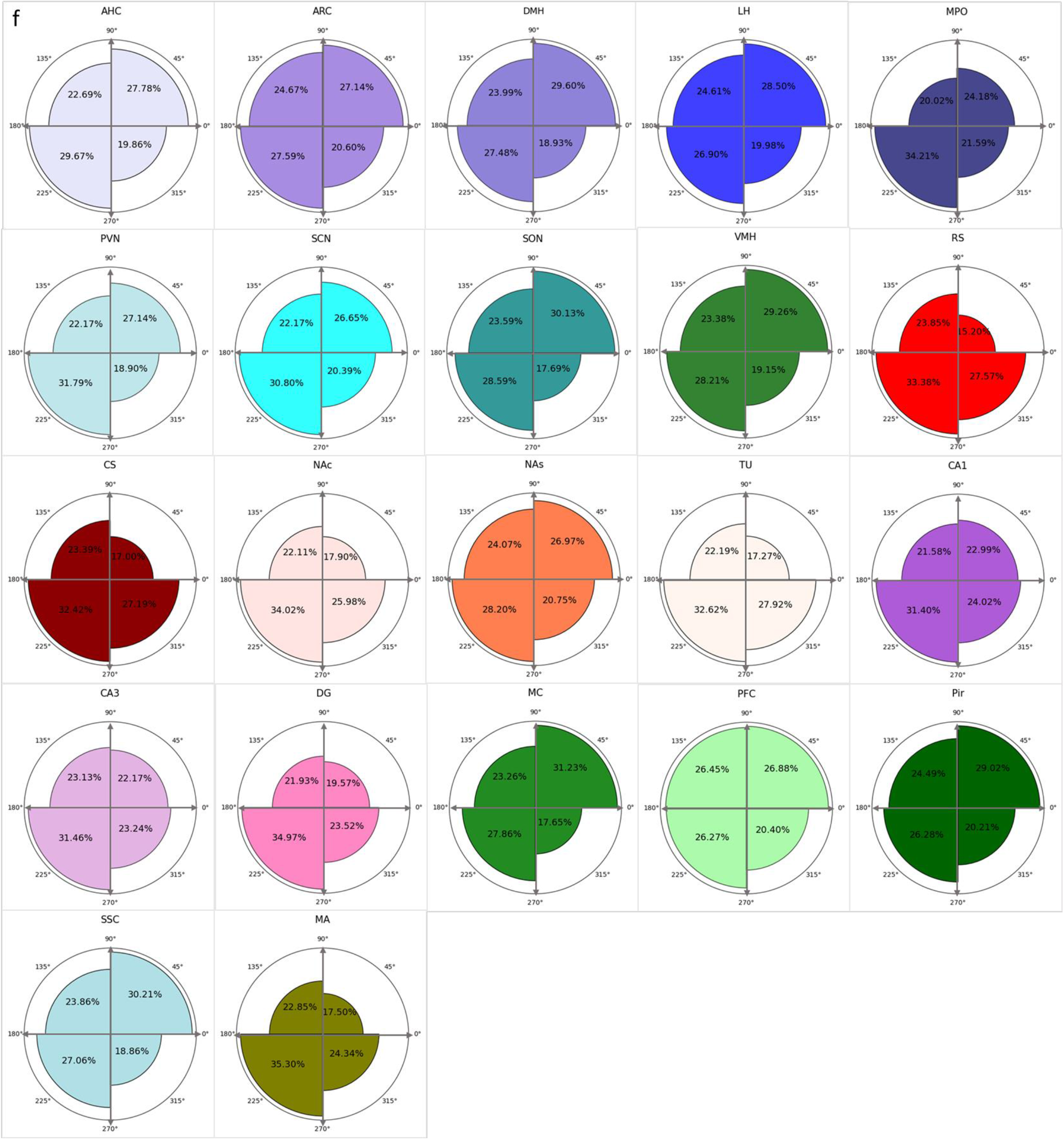
Cilia circular angle analysis, based on individual cilia measurements. **(a)** & **(b)** Graphs showing **(a)** Histogram and **(b)** Radial Plot of the frequency distribution of cilia angles in the whole brain from all individual measured cilia. **(c) & (d)** Graphs showing **(c)** Histogram and **(d)** Radial plot of Cilia angle frequency distribution in the 22 brain regions from all individual measured cilia. **(e)** Radial Plot displaying the frequency distribution of cilia circular angles in individual brain regions from all individual measured cilia. **(f)** Rose diagram representing the percentage of cilia circular angles in the four polar coordinates of cilia angle distribution.

Calculating the average circular cilia angle of individual sections, based on the cilia circular angle means in each brain section, revealed a distinct trend: the piriform cortex (Pir) displayed the highest circular mean cilia angle reaching 267°, whereas the motor cortex (MC) exhibited the lowest mean circular cilia angle measured at 219° (Fig. 4a,b). The distribution of cilia angles favored the third quadrant (180°-270°), particularly around 250° relative to the horizontal line in coronal sections (Fig. 4c-e).

**Figure 4:**
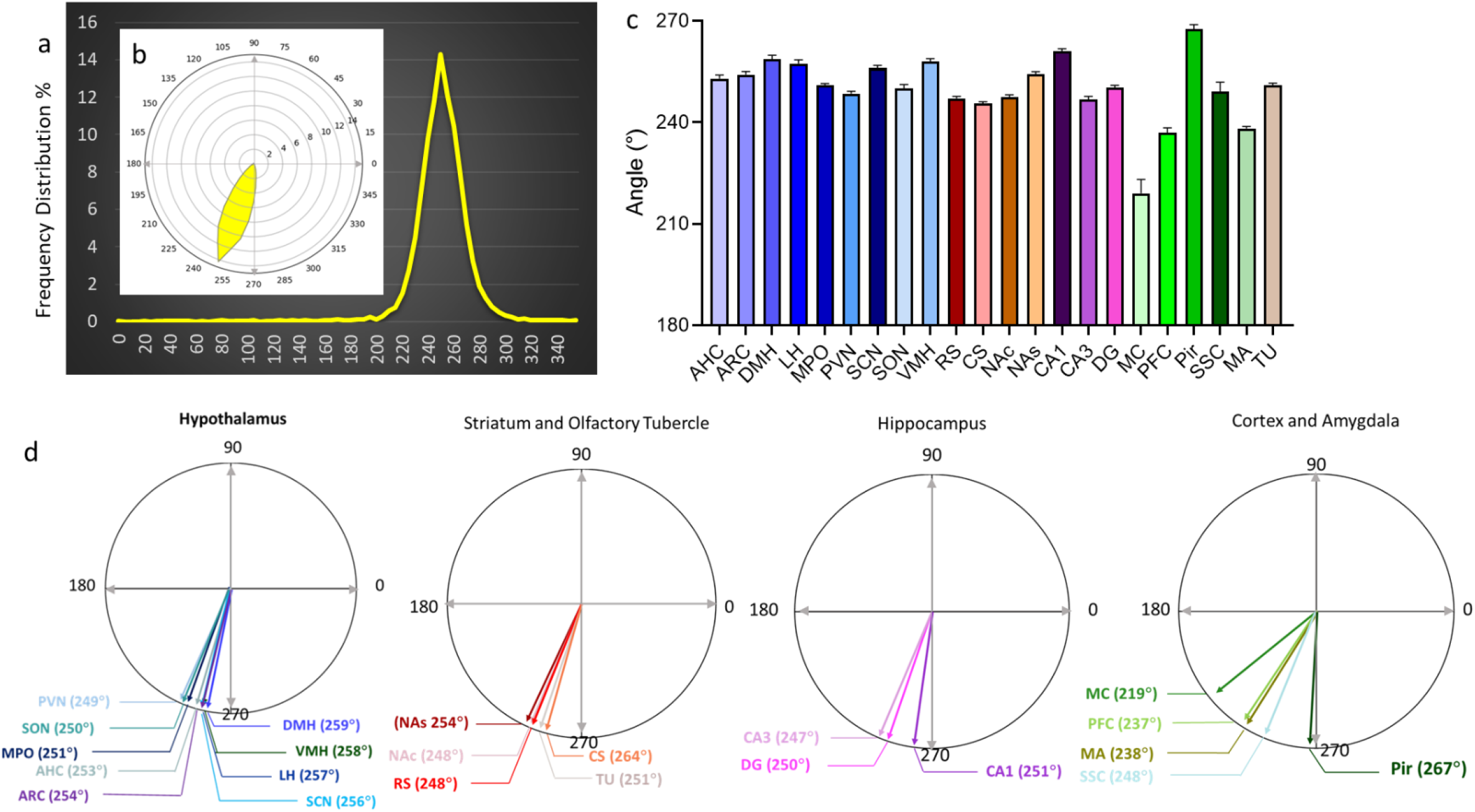
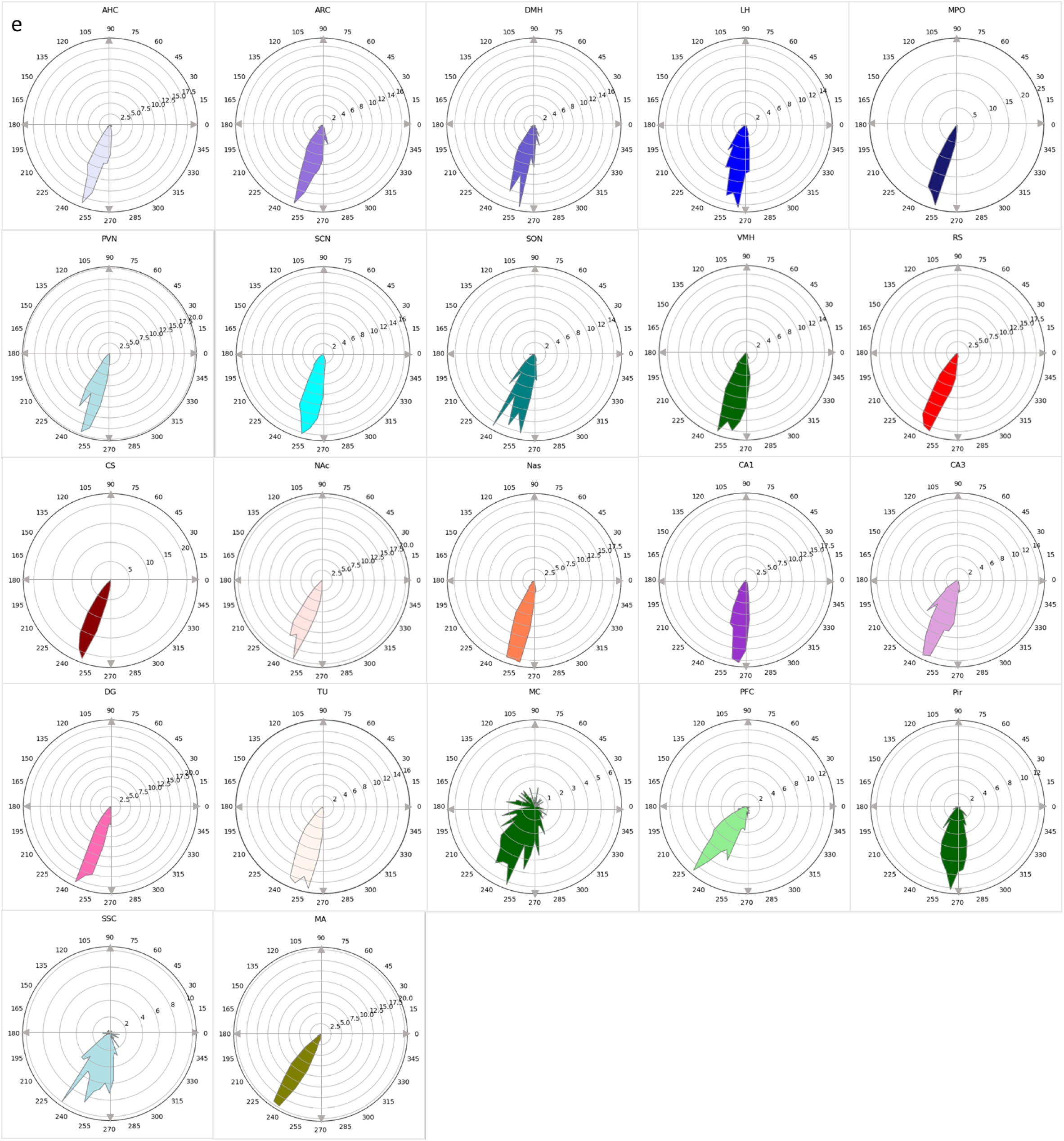
Cilia circular angle analysis, based on sections’ cilia circular angle means. **(a) & (b)** Graphs displaying **(a)** Histogram and **(b)** Radial Plot of the frequency distribution of cilia circular angles’ mean from the mean of all region means in the whole brain. **(c)** Histogram plots of the means±SE of cilia circular angles’ in the 22 brain regions. The average angle is determined based on the cilia circular angle means in each brain section. **(d)** Radial plots of the means of cilia circular angles’ in the 22 brain regions, which are grouped into major categories including, the Hypothalamus, Hippocampus, Cortices, and Striatum. The average angle is determined based on the cilia circular angle means in each brain section. **(e)** Radial Plot showing the frequency distribution of cilia angles from the mean of the section circular means in the 22 individual brain regions.

### Temporal fluctuations in cilia length and orientation across brain regions

We next analyzed the data across the circadian cycle (Fig. 5a-d, S3a-c Table S2). Our analysis revealed complex diurnal fluctuations in both the length and orientation of cilia across brain regions, suggesting a complex temporal regulation of cilia characteristics across different brain areas (Fig. 5a-d, S3a,c). Apart from the AHC and CS, all other brain regions displayed time-dependent variations in cilia length. An examination of the timing for maximal cilia length revealed distinct patterns during light and dark phases (Fig 5a,d, S3a,c). Specifically, during the light phase, maximal cilia length was observed in the MA at 0 and 10 ZT, LH and RS at 2 ZT, CS at 4 ZT, and TU, AHC, CA3, and NAc at 8 ZT. In contrast, throughout the dark phase, peak cilia length was evident in the ARC and DG at 14 ZT, and NAs and PFC at 18 ZT (Fig. 5a,d, S3a). Interestingly, both the DMH and VMH showed a distinct bimodal pattern of maximal cilia length during both the light and dark phases, specifically at 6 and 18 ZT. On the other hand, the SSC presented a different pattern, reaching peak cilia lengths within the intervals of 6-10 ZT and 14-18 ZT (Fig. 5a,d, S3a).

**Figure 5:**
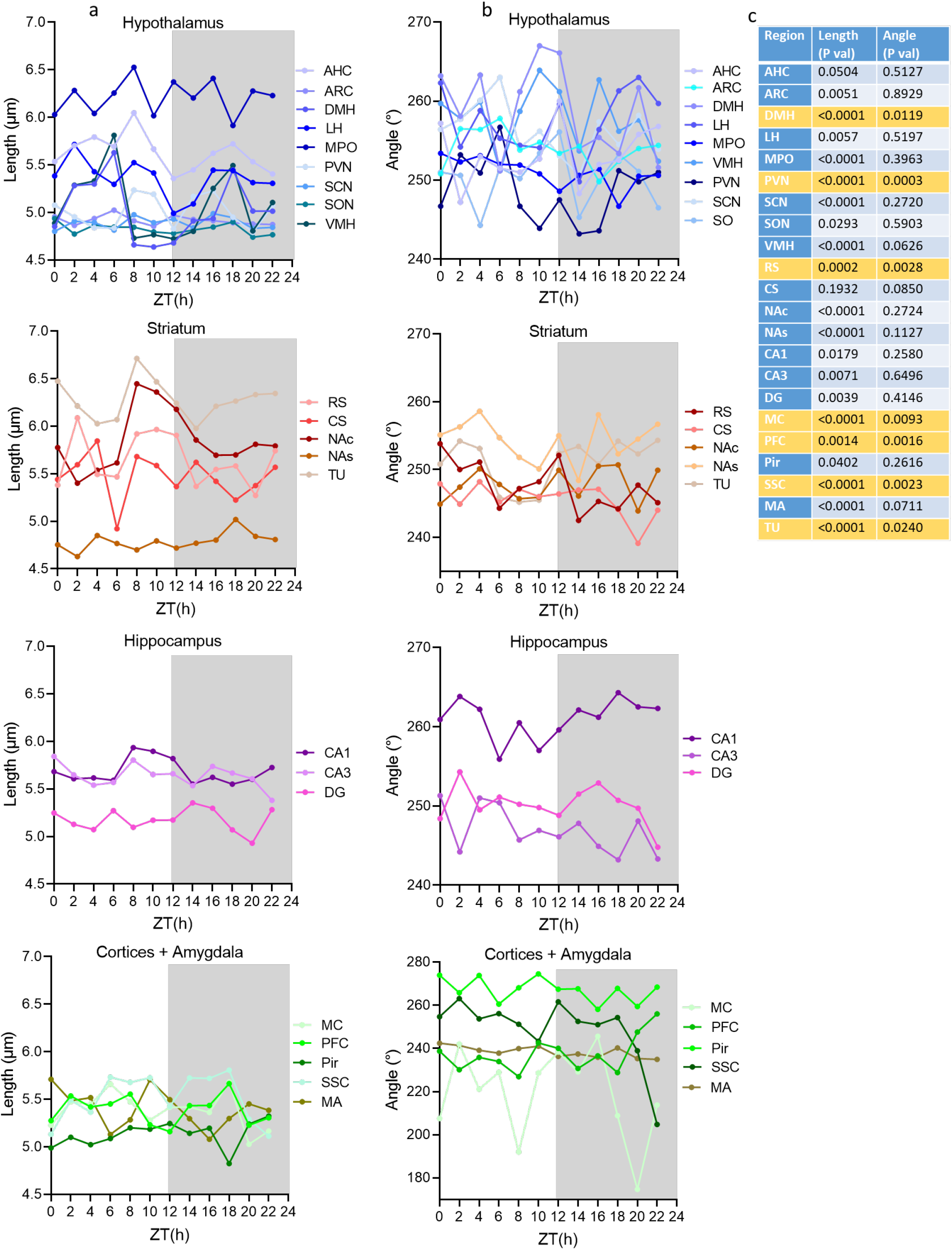
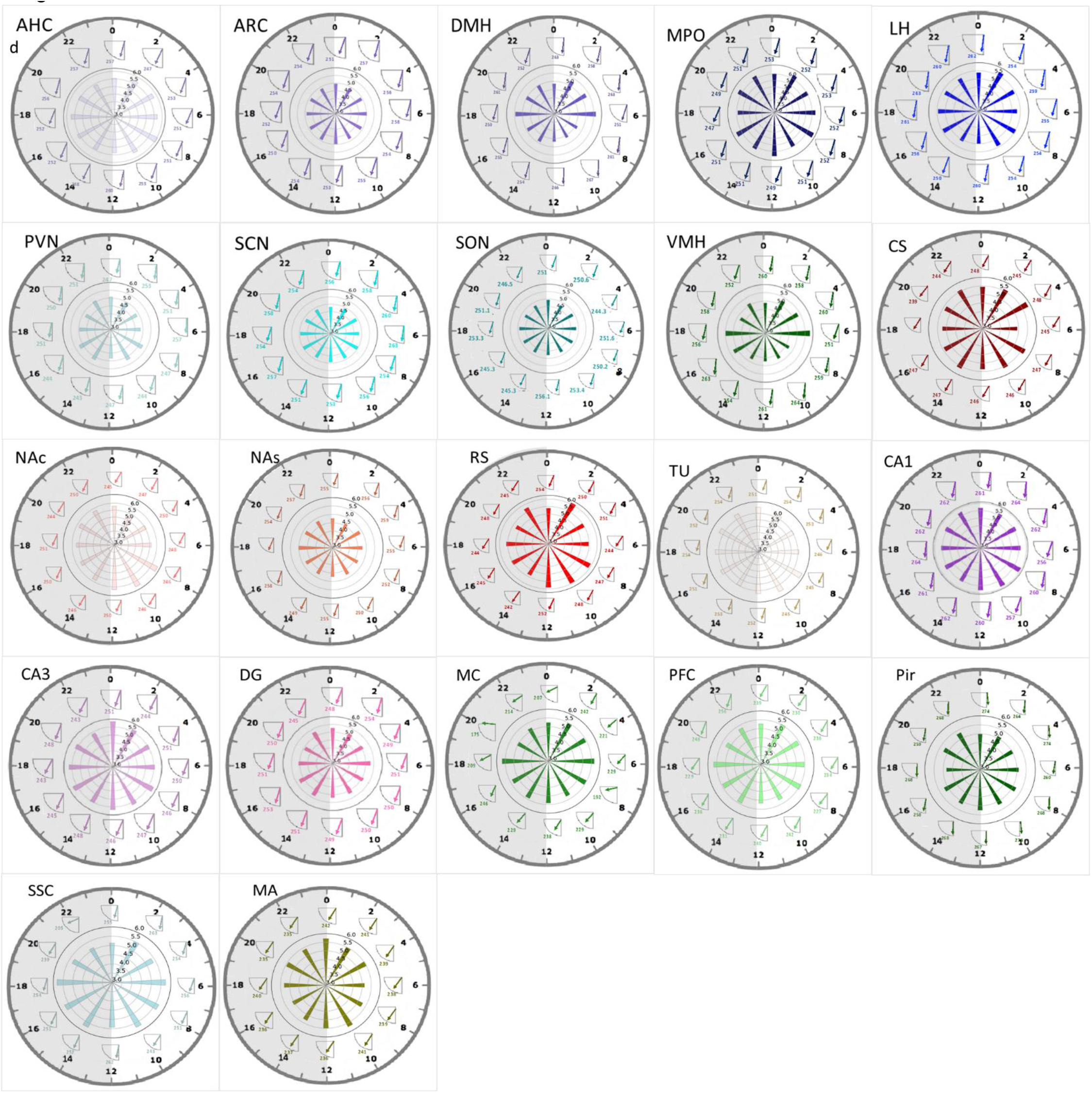
Fluctuation of cilia length and cilia circular angle across 24-hours’ time. **(a & b)** Diurnal fluctuations of cilia length **(a)** and angle **(b)**: The plots show the average cilia length and angle over the course of 24-hours day in different brain regions. The brain regions are grouped into major categories including, the Hypothalamus, Hippocampus, Cortices, and Striatum. The average length and angle are determined based on the measurements’ means in each brain section; Zeitgeber time (ZT). One-way ANOVA test was used to compare the means of lengths and angles at different time points. *P* values were calculated using the False Discovery Rate (FDR) correction, employing the Two-stage linear step-up procedure of Benjamini, Krieger, and Yekutieli for multiple comparisons. **(c)** P values of one-way ANOVA test, used to assess the variability in cilia length and angle at different time points (in **a**,**b,d**), P < 0.05: significant changes across the different time points over 24 hour-period. **(d)** Diurnal fluctuations of cilia length and angle means in individual brain regions across 24-hour time: The plot shows the fluctuations in cilia length and angle means within specific brain regions over a 24-hour period. Light and dark shades represent the light and dark phases. The ZT is displayed on the outer circumference of the large circle. The inner circumference represents the fluctuations of circular angle means at different time points, while the inner circle represents the fluctuations of the cilia length means at different time points.

We also found that cilia angles demonstrated time-dependent changes in 7 brain regions, some of which overlapped with regions exhibiting time-dependent length changes (Fig. 5b-d, S3b,c). These regions included DMH, PVN, RS, MC, PFC, SSC, and TU. All other brain regions did not exhibit significant fluctuations in cilia angles over the course of the day. Two hypothalamic nuclei, specifically the DMH and PVN, exhibited opposite cilia angle patterns: such that during the transition from light to dark (16-20 ZT), DMH exhibited its highest peak cilia angle, whereas PVN exhibited its lowest cilia angle. Conversely, at 6 ZT, this relationship became inverted, with the DMH exhibiting its lowest angle and PVN exhibiting its highest cilia angle (Fig. 5b,d, S3b). Similarly, the PFC exhibited opposite changes to MC and SSC during the transition from dark to light phases, with PFC exhibiting its highest angle peak while the MC and SSC exhibited their lowest cilia angles, respectively, at 22 ZT. MS showed the largest range of changes in cilia angles across 24 hours, from 174° at 22 ZT to 245° at 18 ZT (Fig. 5b,d).

### Cilia length follows circadian rhythm in specific brain regions

The Biocycle program was used to analyze the circadian rhythms of cilia length and angle across various brain regions. Among all the brain regions investigated, we discovered that five regions exhibited significant circadian rhythms in their cilia length. These regions were the ARC, DMH, VMH, NAc, and SSC (Fig. 6). In stark contrast, none of the brain regions exhibited notable circadian rhythms with respect to their cilia angle. In terms of cilia length, we observed that it peaked at different ZT for each of these five regions: ARC peaked at 15 ZT, DMH at 3.8 ZT, VMH at 3.4 ZT, NAc at 11 ZT, and SSC at 12.16 ZT. For cilia angles, both the DMH and VMH showed peak values at 4.6 ZT (Fig. 6).

**Figure 6:**
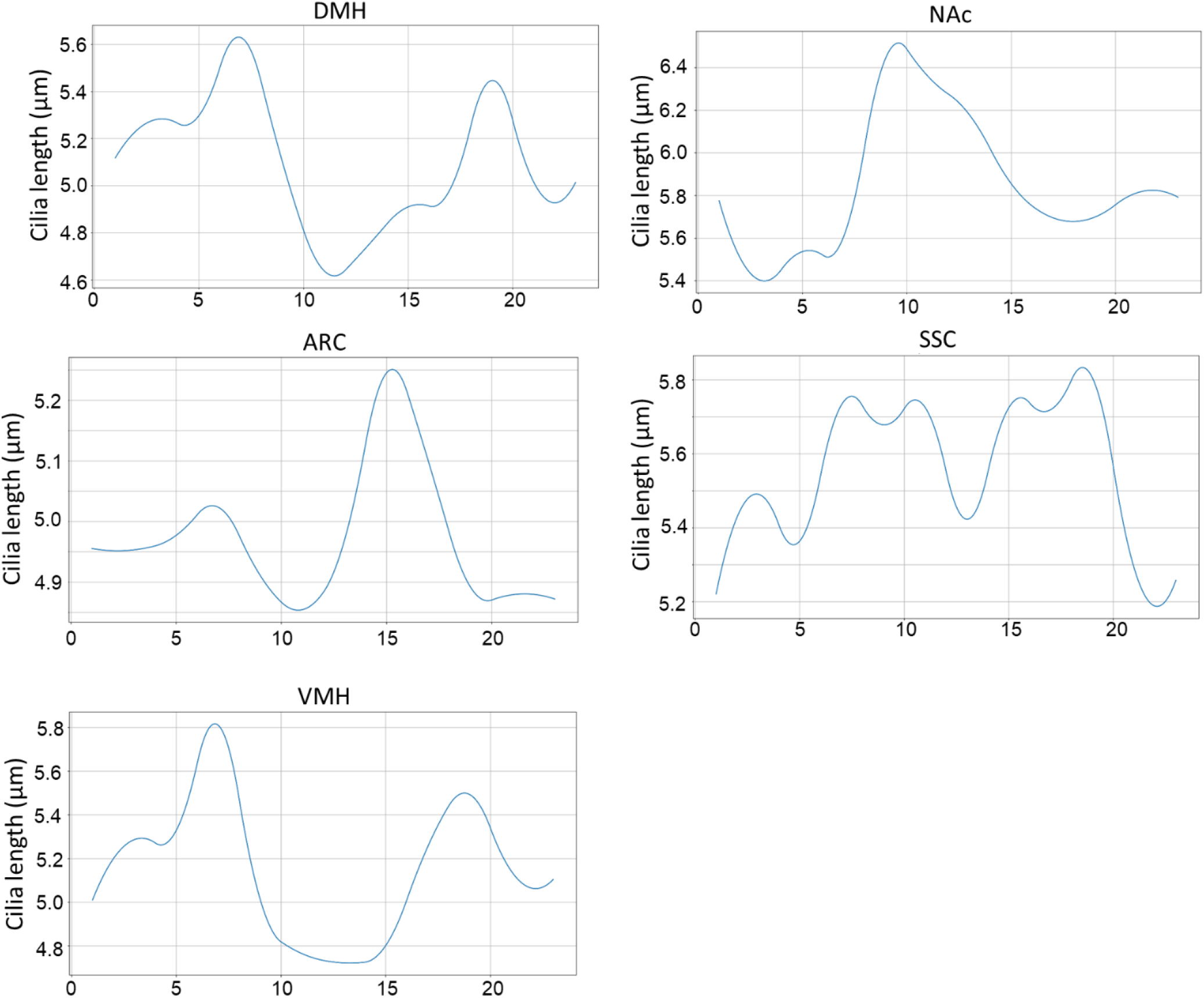
Circadian rhythms of cilia length. BioCycle analysis was used to examine the presence of rhythmic fluctuations in cilia length, and revealed significant circadian patterns of cilia length variations across a 24 hour period in five out of the 22 brain regions tested: ARC DMH, VMH, NAc, and SSC, with p values less than 0.05.

### Correlation of cilia length and circular angle among various brain regions

We next wanted to assess whether fluctuations in cilia length or angle exhibited correlations across brain regions. Our analysis revealed significant correlations within the fluctuations of cilia lengths across different brain regions. Specifically, the length of cilia in DMH and VMH displayed remarkably similar patterns, reflecting a strong correlation with a coefficient (r) of 9.3 (Fig. 7a). Moreover, the lengths of cilia in the PFC demonstrated positive correlations with those in the DMH, VMH, LH, AHC, and MC. In turn, the MC exhibited positive correlations with the DMH, VMH, and SSC. In contrast, cilia lengths in the CA1 region showed negative correlations with the DMH and VMH, but positively correlated with the RS, NAc, and DG. Further negative correlations were found between cilia lengths in the NAc and those in the DMH and VMH, and positive correlation between the TU and the NAc. Lastly, while cilia lengths in the ARC were negatively correlated with those in the TU and positively correlated with the DG, cilia lengths in the LH displayed a positive correlation with those in the AHC (Fig. 7a).

**Figure 7:**
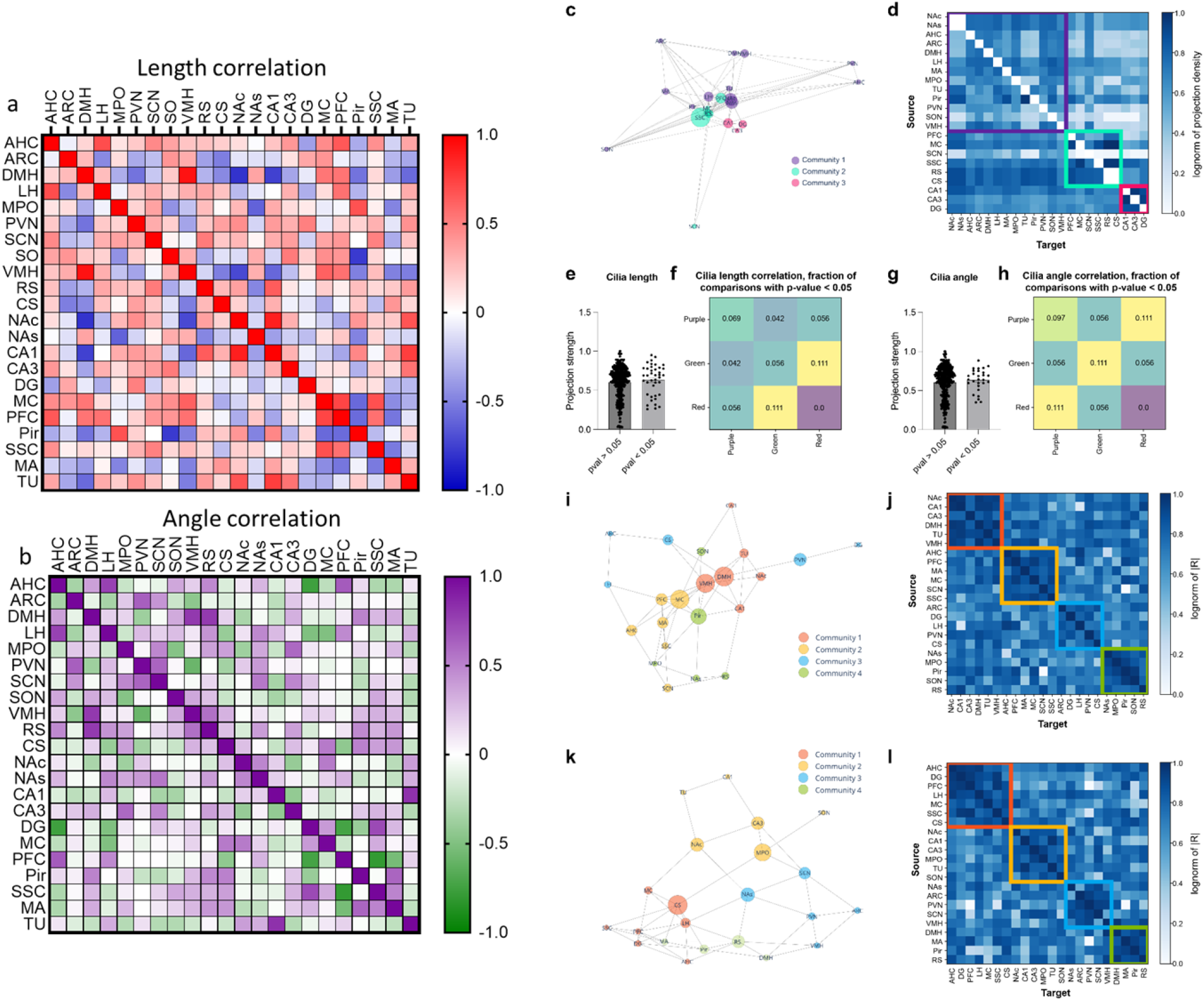
Correlation of cilia length and circular angle among various brain regions and relationship between cilia length/angle and network connectivity. **(a)** Heatmap showing the correlation of cilia length (based on the mean of the section means) among different brain regions. The correlation coefficient ranges from 1 (indicating the most positive correlation, red) to -1 (indicating the most negative correlation, blue). Positive correlation implies that the cilia lengths in the two regions are on the longer and shorter side simultaneously. Negative correlation indicates that the two regions have opposite cilia length sizes. **(b)** Heatmap displaying the correlation of cilia angle (based on the mean of the section circular means) among different brain regions. The correlation coefficient ranges from 1 (indicating the most positive correlation, purple) to -1 (indicating the most negative correlation, green). **(c)** Anatomical network of the brain regions in our study. The network is embedded in 2D space using a spring embedding, in which highly connected nodes are pulled together as by a spring. Node size represents betweenness, a measurement of centrality based on the number of shortest paths between nodes that a particular node shows up on. Nodes are colored according to their community assignment, as determined by the Leiden algorithm. **(d)** Correlogram indicating the strength of input-output connections between all pairs of regions in the dataset. The color bar indicates the log-normalized projection density between any two regions. Regions are grouped according to their community membership. **(e)** Bar graph comparing projection strength between regions significantly correlated with cilia length vs those not correlated. Projection strength equals the log normalization of normalized (by injection volume) projection density. **(f)** Fraction of region pairs significantly correlated by cilia length among all regions pairs between two communities. For example, a fraction of 0.111 means that 11.1% of all region pairs between the red and green communities are correlated by cilia length. **(g)** Bar graph comparing projection strength between regions significantly correlated with cilia angle vs those not correlated. **(h)** Fraction of region pairs significantly correlated by cilia angle among all regions pairs between two communities. **(i)** Cilia length correlation network. The spring embedding puts regions more highly correlated closer together. Betweenness and community membership are shown as node size and color, respectively. **(j)** Cilia length correlogram shows the groupings of different regions by community, as well as their correlation patterns. **(k)** Cilia angle correlation network. Betweenness and community membership are shown as node size and color, respectively. **(l)** Cilia angle correlogram shows the groupings of different regions by community, as well as their correlation patterns.

Similar to cilia length, our analysis also revealed substantial correlations amidst the fluctuations of cilia angles throughout numerous brain regions. The patterns of cilia angle in the DMH and VMH demonstrated a strong correlation, with a correlation coefficient of 0.79 (Fig. 7b). A positive correlation was found amongst the temporal changes in cilia angles across three other pairs of hypothalamic regions, namely the AHC-LH, ARC-PVN, PVN-SCN, and negative correlation was found between PVN and VMH. Further, the cilia angle in the PFC, exhibited positive correlations with the AHC and negative correlations with SSC and DG. In turn, DG cilia angles fluctuations correlated negatively with AHC, and positively with SSC. Cilia angles in the DMH also showed a positive correlation with those in the RS. Notably, cilia angle in the CA1 region correlated positively with the SSC and negatively with the TU, and CA3 showed a positive correlation only with the MPO. Finally, the changes in cilia angle in the Pir exhibited a positive correlation with those in the MA (Fig. 7b).

We next wanted to test the hypothesis that more highly correlated brain regions, either in cilia length or angle, may be preferentially interconnected. To do this, we created a weighted, directional anatomical connectivity matrix using publicly-available projection data from the Allen Mouse Brain Connectivity Atlas (Fig. 7c). We then used a Louvain community detection algorithm to assign regions into “communities”, defined by their holistic connectivity relationships with other nodes in the network. Based on these data, our unsupervised approach identified three unique clusters of brain regions, which we color-coded. The first and smallest community (“red”) contained the three hippocampal regions DG, CA1, and CA3. The second community (“green”) contained cortical regions, the striatum, and somewhat surprisingly, the SCN. The third community (“purple”) contained all of the other regions, which overall had less strongly-defined interconnectivity relationships (Fig. 7d).

Based on these data, we then asked if regions that had strong length correlations (p < 0.05) were more likely to have stronger interconnectivity than regions with weak length correlations (p > 0.05). We found no significant difference in normalized projection strength between pairs of regions correlated with cilia length versus those not correlated (Fig. 7e), indicating that projection strength between regions does not influence cilia length. To further test if inclusion within specific communities may influence correlation patterns, we examined all pairs of regions within and across different clusters and identified a bias towards regions having correlated cilia lengths if they were present within particular communities; this relationship was most prominent between regions in the red and green communities (Fig. 7f). In order to test the significance of this observation, we shuffled p-values 1000 times across all pairs of regions, and examined the fraction of correlated region pairs between sets of communities. We found that the fraction of correlated pairs between red-green communities (0.111) was in the st percentile relative to the random distribution, indicating that cilia lengths were more likely than by chance to be correlated between pairs located in red and green communities (Fig. 7f).

We next tested if regions that had strong angle correlations (p < 0.05) were more likely to have stronger interconnectivity than regions with weak angle correlations (p > 0.05). As with cilia length, we did not identify a significant difference in projection strength between pairs of regions correlated with cilia angle versus those not correlated (Fig. 7g). We then assessed the relationship between community identity and cilia angle correlation, and found that the fraction of correlated pairs within the purple community (0.107) was 7.4th percentile relative to the random distribution (Fig. 7h). These data together indicate that while the connectivity strength between directly connected pairs did not appear to relate to cilia length or angle, the presence within particular communities did appear to relate to both cilia length and angle, indicating that the network architecture of the brain exhibits some relationship with cilia length and angle correlations across regions.

Given the lack of clear relationship between direct anatomical connectivity and cilia length or angle correlations, we next applied the network approach on the cilia length and angle correlation data directly. In order to do this, we built correlation networks using a log scaled R value that yielded a nicer distribution for edge weights, and again used the Louvain community detection algorithm to assign communities (Fig. 7i-l). These results yielded four communities that did not distinctly track by anatomical connectivity, as evidenced by a different regional distribution across communities. Interestingly, we observed a number of regions that cluster together in both cilia length and angle correlation networks. Notably, NAc, CA1, CA3, TU cluster together for both, as do AHC, PFC, MC, and SSC (Fig. 7i-l). The NAc and TU are both striatal regions while CA1 and CA3 are both hippocampal, and PFC, MC, and SSC are all cortical regions; however, the exact relationship between these sets of regions that co-cluster is not immediately apparent, again suggesting that factors other than direct connectivity likely influence the non-random correlations in cilia length and angle across regions.

### Correlation between circadian physiological variables and diurnal fluctuation patterns of cilia length and angle

Lastly, we analyzed the correlation between the diurnal fluctuation patterns of cilia length and angle and various parameters known to follow similar 24-hour rhythms (Fig. 8,a,b, S4a). In four hypothalamic regions, the 24-hour fluctuation patterns of cilia length exhibited a positive correlation with parameters such as melatonin levels (DMH), and negative correlations with locomotor activity (AHC and LH), food intake (DMH and VMH), and body temperature (AHC and LH) and displayed a positive correlation with NREM sleep episode duration (LH) (Fig. 8a, S4a). In the striatal structures, we found a negative correlation between cilia length’s 24-hour pattern and temperature (AHC), glutathione (GSH) levels (RS), melatonin (CS), glutathione S-transferase (GSH ST) activity (CS), and NREM EEG delta sleep (NAc). On the other hand, there was a positive correlation between cilia length and REM EEG theta sleep (NAs). The 24-hour fluctuation patterns of cilia length within CA1 and PFC displayed a negative correlation with NREM EEG delta sleep and food intake, respectively (Fig. 8a, S4a).

**Figure 8:**
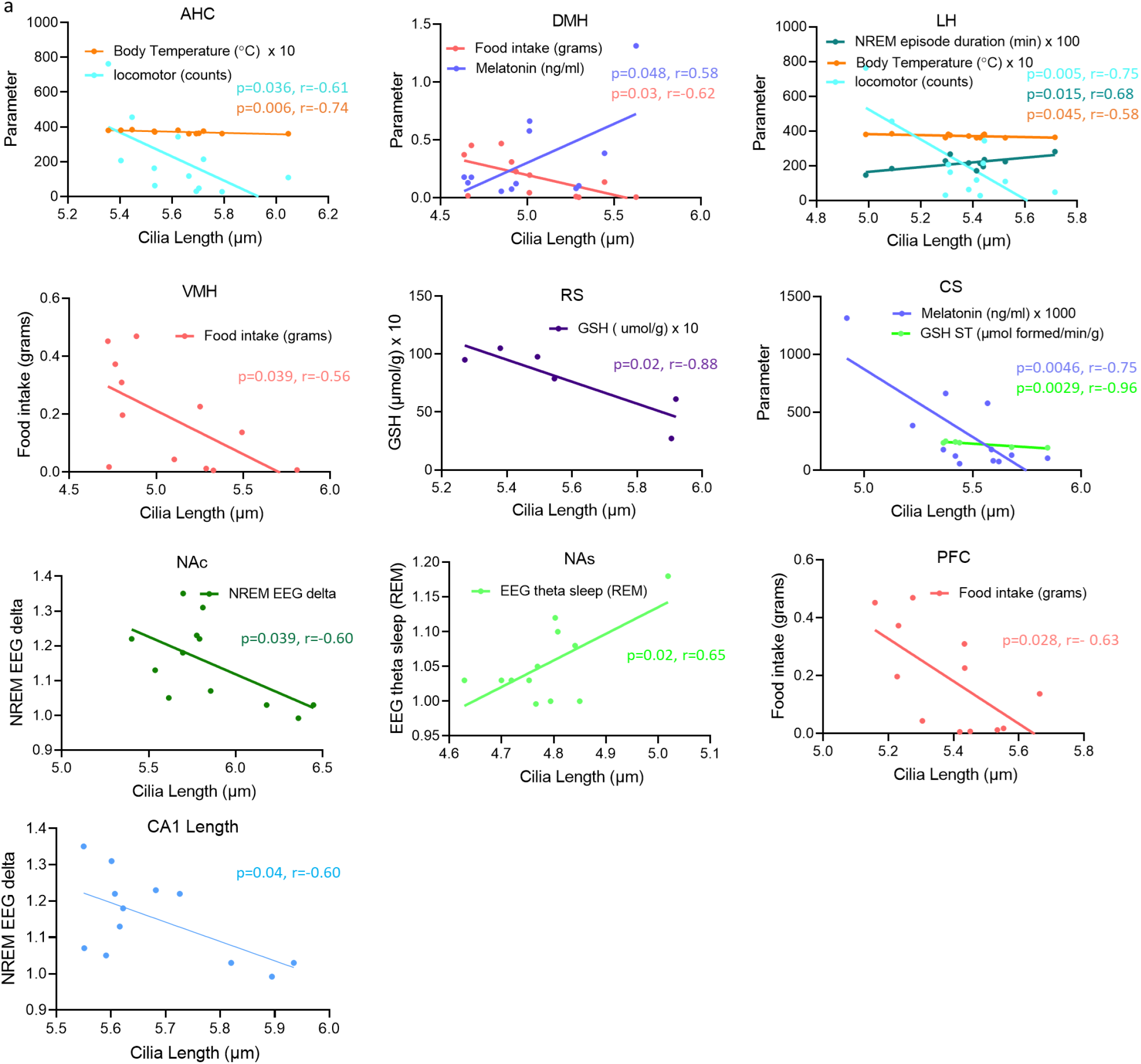
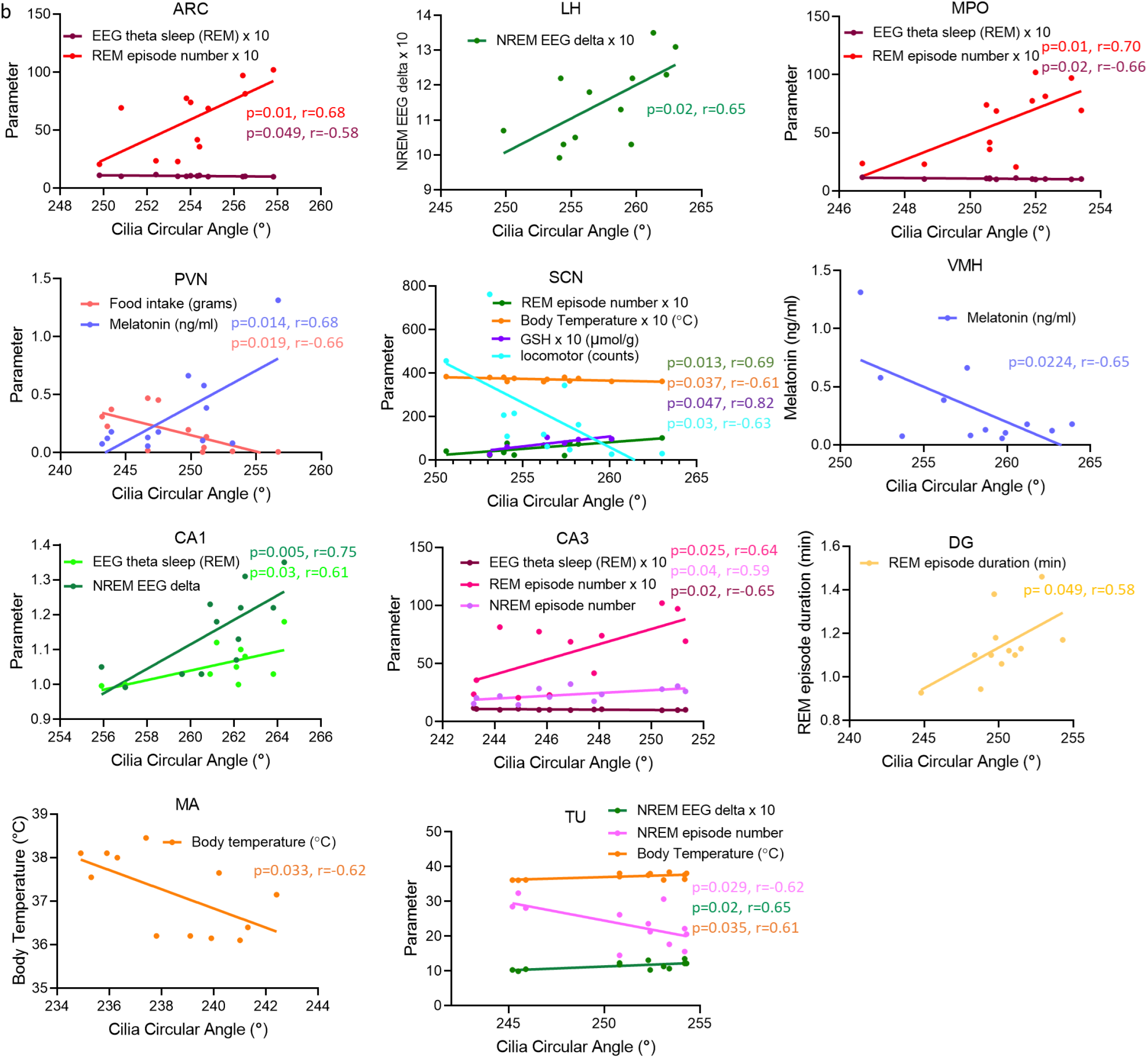
Linear regression relationship between rhythmic variables and Cilia length/angle. Scatter plots and fitted linear regression lines displaying the relationship between cilia **(a)** length and **(b)** circular angle and various circadian parameters. Each plot represents the correlation between rhythmic variables and Cilia length/angle in one region. Only the parameters that show significant correlations (p < 0.05) with cilia length/ circular angle are included in the plots. The correlation coefficient (r) reflects the strength and direction of the correlation: positive r indicates a positive slope, and negative r indicates a negative slope.

We also found that the diurnal variations in cilia angle within six hypothalamic regions displayed correlations with numerous parameters. These included positive correlations with REM episode count (ARC, MPO, and SCN), NREM EEG delta (LH), melatonin (PVN), and GSH (SCN), along with negative correlations with EEG theta during REM sleep (ARC and MPO), locomotor activity and body temperature (SCN), food intake (PVN), and melatonin (VMH) (Fig. 8b, S4b). Furthermore, cilia angle fluctuations over a 24-hour period in hippocampal structures exhibited positive correlations with EEG theta during REM sleep and NREM EEG delta sleep (CA1), REM episode count and NREM episode count (CA3), REM episode duration (DG), while exhibiting a negative correlation with EEG theta during REM sleep (CA3) (Fig. 8b, S4b). Finally, cilia angle variations across a 24-hour cycle displayed negative correlations with temperature in the MA and positive correlations in the TU, while showing a positive correlation with NREM EEG delta and a negative correlation with NREM episode number in the TU (Fig. 8b, S4b).

## Discussion

In our study, we examined the spatiotemporal patterns of cilia length and orientation across 22 mouse brain regions. Our findings revealed substantial variation in cilia length and direction across these regions, demonstrating the non-randomness of their orientation, and highlighted distinct spatiotemporal and circadian patterns in certain regions.

We developed automated image analysis algorithms, which allowed us to conduct a high-throughput analysis of the spatiotemporal variability and circadian rhythms of primary cilia length and orientation across mouse brain regions. These algorithms enabled us to examine over 10 million individual cilia, generating over 240 million data-points, and yielding the largest spatial and temporal atlas of cilia metrics. We found that cilia length varied across distinct brain regions, with the shortest average cilia length observed in the NAs and the longest in the TU. Likewise, cilia angles differed across brain areas, with the smallest average angle observed in the MC and the largest angle in the Pir.

The noticeable variation in cilia length among different brain regions suggests tailored functional requirements for each area. Longer cilia might offer increased surface area and accommodate more receptors, thereby enhancing sensitivity to extracellular signaling molecules and promoting more effective signal transduction between cells.

The finding that cilia display distinct orientation patterns across various brain regions presents a novel discovery. The distinct distribution at 45° intervals indicates that the orientation of cilia within the brain is not random but follows distinct patterns. Furthermore, the prevalent orientation towards the medial-ventral quadrants in most of the examined brain regions affirms this notion of a non-random distribution. Interestingly, cilia orientation appears to exhibit a remarkable congruity within anatomically or functionally interconnected structures. This is observed, for instance, in striatal structures RS, CS, and NAc, as well as hippocampal structures, including CA1, CA3, and DG, which all show a distinct angle distribution primarily towards 180° and/or 270°. Interestingly, the TU, also categorized as a striatal structure, mirrors this orientation pattern of the striatum. Further, the medial amygdala (MA), known for its substantial neural connections with the striatum, exhibits a similar cilia orientation pattern (towards 180° and/or 270°). In contrast, the hypothalamic nuclei, cortices, and NAs display a more diverse cilia orientation pattern within the 45° interval range. Despite this variability, the majority still lean towards the medial-ventral quadrant or along the axis from 45° to 225°—the minor diagonal in a Cartesian coordinate system. Our findings suggest that shared connectivity, and thus shared functional or anatomical properties, may influence the pattern of cilia orientation, and reinforce the notion that the distribution of cilia orientation across the brain is not random but rather is linked to the distinctive functional or anatomical characteristics of these interconnected regions.

While not clearly related to anatomical connectivity patterns, these distinctive cilia orientation patterns across various brain regions may relate to their specific locations or proximities to the associated ventricles and their particular functions. Cilia orientation patterns across different brain regions are likely influenced by several interconnected factors. For example, the NAc, TU, CA1, and CA3 all associate within the same community based on cilia length or angle correlations (Fig. 7), and all are correlated with some aspect of sleep (Fig. 8). In addition, the location of a brain region in relation to specific ventricles might be an important factor. For instance, cilia present in areas like the striatum, TU, and hippocampus predominantly orient towards the lateral ventricles. Conversely, those in the NAc and TU have a stronger connection to the third ventricle. Cilia in hypothalamic nuclei also seem to align more closely with the third ventricle, while those in the cortices orient towards both the ventricles and the subarachnoid space. Additionally, the unique structure and function of each brain region, alongside the chemical gradients and cerebrospinal fluid flow from associated ventricles, may further shape these orientation patterns. Given their directional sensitivity, cilia orientations could have significant implications for how brain regions respond to environmental cues.

Our comprehensive analysis revealed that both cilia length and angle in numerous brain regions exhibit time-of-day fluctuations. Except for the CS and AHC (*P*=0.05), all the regions tested, displayed time-dependent fluctuations in cilia length, and seven of these regions also showed corresponding fluctuations in cilia orientation. The observed changes in cilia length across the light and dark phases in different brain regions may correspond to varying sensory or signal transduction requirements during these periods. For example, an increase in cilia length may enhance the detection or transmission of specific signals. In the context of the hypothalamus, which plays a crucial role in maintaining circadian rhythm, the bimodal pattern of maximal cilia length observed in DMH and VMH might be associated with their role in controlling distinct behavioral or physiological processes that peak at different times of day. Thus, our findings underscore the pivotal role of cilia dynamics in directing an extensive range of time-dependent physiological functions such as sleep-wake cycles, feeding behavior, regulation of body temperature, hormone secretion, and stress responses [35-42]. The finding that cilia angles also demonstrate time-dependent changes in certain brain regions is especially interesting. This suggests that in addition to the varying lengths, the direction of signal sensing or transmission by cilia might also shift over the course of a day. The observed inverted relationship between the DMH and PVN, and between PFC, MC, and SSC, could represent complementary roles of these regions in processing or transmitting different signals during the light and dark phases. Temporal changes in cilia length and orientation could impact the sensitivity of receptors localized on cilia towards their ligands, or influence the efficiency of signal transmission, ultimately affecting the brain’s ability to regulate these time-sensitive processes.

Our Biocycle analysis uncovers significant circadian rhythms in cilia length spanning across five pivotal brain regions, thereby unfolding the complex landscape of ciliary dynamics and its profound implications on circadian physiology. The NAc, a vital hub for reward and motivation, presents a distinct rhythmicity in cilia length, which calls into question whether the physical changes in cilia length may be a contributing factor to the timing of reward-seeking behaviors. Cilia have been shown to play distinct roles in the acute and long-term responses to psychoactive drugs [62], which potentially underpins circadian variations in reward-seeking behavior. Interestingly, despite the prominent role of the SCN in the orchestration of circadian rhythms within the brain, our study did not detect any circadian regulation of cilia length or angle in this region. While we observed variability in SCN cilia length across the 24-hour period, these changes did not appear to follow a circadian rhythm. This is notably different from a recent study, which showed a pronounced circadian pattern in SCN cilia length with a significant 5-fold difference throughout the day [63]. The reasons underlying this discrepancy are unclear. In our study, we used a large-scale unbiased comprehensive approach to assess cilia length and angles across numerous brain regions, using Swiss Webster mice. This approach differs from the previous study which focused exclusively on the SCN, collecting data every 6 hours over a 48-hour period. Furthermore, we utilized the BioCycle program for our analyses, and we collected our data at two-hour intervals within a 24-hour span. The ARC, DMH, and VMH - critical hypothalamic regions known for orchestrating energy balance, feeding behavior, and sleep-wake cycles [64-68], also display significant circadian rhythms in cilia length. This points towards the intriguing possibility that these rhythmic changes in cilia length may serve as a physical manifestation of their functional roles in these circadian processes, adding a structural dimension to our understanding of these critical regulatory pathways. The SSC’s observed circadian rhythms in cilia length is interesting in light of previous studies showing rhythmic synaptic changes in the SSC [69-72], pointing at cilia’s role in modulating sensory processing throughout the day-night cycle. These findings collectively emphasize the potential involvement of cilia in modulating various circadian behaviors and physiological processes, and may offer valuable insights into the mechanisms underlying circadian rhythms.

The spatiotemporal patterns we observed in cilia length and angles across the 22 studied brain regions highlight a multifaceted and diverse landscape of cilia dynamics. Several of the hypothalamic nuclei investigated in our study play integral roles in controlling a variety of time-sensitive processes, such as REM and NREM sleep, food intake, energy metabolism, hormone secretion, and stress responses. Our observations suggest that fluctuations in cilia length and orientation, exhibiting peaks during both light and dark phases, could play a crucial role in fine-tuning various physiological processes. These peaks in cilia length may mirror the functional needs of specific regions at different times of the day. Intriguingly, we noted the shortest cilia lengths in hypothalamic nuclei, with the exception of the SCN and MPO, occurring during the transition from dark to light phase or light to dark phase. This observation may hint at a potential role for cilia in detecting environmental changes and consequently modulating the output of hypothalamic nuclei. In particular, hypothalamic regions such as the DMH, LH, anterior hypothalamus, ARC, and VMH displayed peaks in cilia length during both the dark and light phases. This could indicate their involvement in processes requiring coordinated regulation across day and night cycles, such as feeding behavior, body temperature regulation, or sleep-wake transitions. Furthermore, it is worth noting that cerebrospinal fluid (CSF) composition and flow dynamics follow time-dependent patterns [73]. This further highlights the potential relationship between cilia dynamics and the regulation of various physiological functions demonstrating time-of-day dependencies. Cilia are known to house numerous receptors, including G-protein coupled receptors (GPCRs) and ion channels, which serve a critical role in detecting extracellular signals and translating them into intracellular responses. Notably, several of these cilia-localized receptors play a role in regulating various circadian functions. For example, receptors such as melanocortin 4 receptor (MC4R), melanin-concentrating hormone receptor 1 (MCHR1), neuropeptide Y receptors (NPY2R, NPY5R), somatostatin receptor 3 (SSTR3), serotonin receptor 6 (5HT6), and dopamine receptor 1 (D1) are integral to the regulation of sleep-wake cycles and feeding behavior [37, 43-47]. Thus, the time-of-day dependent fluctuations in cilia length and orientation could potentially modulate the sensitivity of these receptors to their respective ligands, thereby impacting signal transduction and influencing the brain’s control over these processes.

The correlations we observed between specific brain regions in their cilia length and angles raise intriguing possibilities of common regulatory mechanisms, functional connections, or synchronized responses to environmental stimuli. The 22 brain regions we examined can be thought of as clusters or functional networks, based on their similarities in cilia properties and the functions they regulate. For instance, the cilia length and angle in the DMH and VMH exhibited remarkably similar patterns, which could be indicative of a shared functional relationship between these two areas. Similarly, a positive correlation in cilia length between the PFC and MC might suggest they share a regulatory mechanism or functional connection, possibly related to higher cognitive and motor processing. In addition, the positive correlation observed between cilia length in the ARC and DG could hint at a functional relationship between these regions in regulating feeding behavior and processing spatial or contextual information. The findings that cilia length and angle within different brain structures correlate with key physiological parameters such as melatonin levels, locomotor activity, food intake, body temperature, and sleep parameters offers a tempting evidence for cilia role in the brain’s circadian machinery. For instance, the positive correlation between cilia length in the DMH and melatonin levels suggest a role for cilia in sensing or responding to this key hormone, which is known to regulate sleep-wake cycles and other circadian rhythms. Similarly, the negative correlation between cilia length and locomotor activity or food intake in regions like AHC, LH, and VMH might suggest a potential role for cilia in modulating these behaviors.

It is particularly noteworthy that changes in cilia angle, like those in cilia length, are correlated with several physiological parameters, suggesting that the orientation of cilia may also be dynamically regulated according to the body’s circadian rhythms. This is evident from the positive correlations found between cilia angle variations and factors such as REM episode count, NREM EEG delta, melatonin, and GSH in multiple hypothalamic regions. Furthermore, the correlations observed within hippocampal structures, particularly with sleep parameters, suggest that cilia orientation may also be involved in cognitive processes such as learning and memory, which are known to be influenced by sleep [74, 75]. Lastly, the correlation of cilia angle variations with temperature in MA and TU and sleep parameters in the TU suggests that cilia may have a function in sensory or signaling mechanisms involved in the perception or regulation of body temperature and sleep patterns.

Investigating the molecular and cellular mechanisms underpinning these observed correlations could offer valuable insights into the role cilia play in integrating and coordinating functions across different brain regions.

## Conclusion

In conclusion, our comprehensive investigation into the spatiotemporal dynamics of cilia length and orientation across distinct brain regions reveals a multifaceted landscape of cilia properties. The significant regional variations, the time-of-day fluctuations, and the circadian rhythms observed in certain regions underscore the diverse roles that cilia could play in the brain. Intriguing correlations between specific brain regions suggest potential shared regulatory mechanisms or functional connections. While the precise functional implications of these dynamic cilia properties are yet to be fully understood, our findings highlight the importance of cilia in responding to environmental cues and regulating time-dependent physiological processes.

The physiological relevance of these diurnal variations in cilia length and orientation is an open question. Are these changes necessary for maintaining circadian rhythms, processing different types of sensory input, or carrying out specific neuronal functions at different times of day? Moreover, do these changes contribute to the pathophysiology of neurological disorders that are known to have a diurnal variation in their symptoms? Future research might consider these questions to better understand the functional implications of these novel findings.

## Data Availability

All codes used in the analysis and original data are available in the supplementary materials. Data that support the findings of this study are available from the corresponding author upon request.

## Author contributions

AA designed the experiments and wrote the manuscript; PB designed the analysis methods, generated the codes for cilia length and angle measurements, and measured these parameters supervised the coding, and edited the manuscript. RM conducted the animal and immunohistochemistry experiments and imaging; SA generated the codes for cilia length and angle measurements and measured these parameters; KC, HW, KL, TC, JL, SV, helped with the immunohistochemistry experiments and imaging. PD and KB conducted the network analysis. SN contributed to the experiment design and manuscript editing.

